# The Quest for a Universal Parkinson’s Transcriptomics Signature is Derailed by Inherent Variability between Patients

**DOI:** 10.1101/2025.08.20.671280

**Authors:** Roy Dayan, Serafima Dubnov, Hagit Turm, Michelle Grunin, Shahar Shohat, Salim T Khoury, Ami Citri, Tamar Harel, David Arkadir

## Abstract

**Objective:** To systematically evaluate the reproducibility and clinical utility of published blood mRNA-based gene signatures for classifying PD from healthy controls, and to uncover the pitfalls that limit their performances.

**Methods:** We validated the classification performance of 13 gene signatures unique for PD (published 2015-2025) using the Parkinson’s Progression Markers Initiative (PPMI) database. We further validated classification performance on data collected in a prospective clinical trial where demographic and clinical parameters were minimized and environmental confounders were strictly controlled. Sources of gene expression variability were studied by mean pairwise distance analysis.

**Results:** Gene overlap between signatures was low (11/411, 2.7%) but statistically significant (*P*<0.001) and enriched with genes involved in lipid metabolism. The majority (10/13) of these signatures retained statistical significance when tested on the PPMI dataset, but their classification performance was modest (median AUC 59.7%), substantially lower than originally reported. Classification performance improved when comparing *GBA1*-associated PD to controls (median AUC 65.4%, *P*=0.006). A prospective trial (16 PD patients, 14 controls) with rigorous environmental standardization did not improve classification accuracy. Inter-individual expression variability is the dominant limiting factor in classifier performance. Variability was not impacted by long-term levodopa therapy.

**Interpretation:** While blood transcriptomic signatures unique to PD are reproducible and may elucidate PD pathophysiology, their broad clinical utility remains severely limited by inherent inter-individual variability. Future research should prioritize biomarkers of divergence rather than the pursuit of a universal PD signature.

## INTRODUCTION

Over the past decades, considerable efforts have been devoted to the identification of biomarkers that could enhance the diagnosis of Parkinson’s disease (PD), facilitate the understanding of its pathophysiology and assess the efficacy of potential disease-modifying therapies.^1^ Blood transcriptomics has emerged as a promising tool in this pursuit, offering a minimally invasive method to capture broad, unbiased molecular information.^2^ Several studies have investigated blood transcriptomics for a unique signature of PD, often using machine learning algorithms to manage high-dimensional data, and have reported success in distinguishing PD patients from healthy controls.^3–6^ Yet, despite the persistence of these efforts and the encouraging results, why have transcriptomic biomarkers not been translated into clinical practice?

Previous studies have highlighted several limitations that have hindered the translation of transcriptomics-based approaches into clinical practice. First, RNA expression exhibits substantial variability, even within homogeneous cell populations.^7^ Second, many transcriptomics-based classification methods and machine learning algorithms have been criticized for their susceptibility to circular reasoning^8^ and overfitting.^9^ Consequently, gene signatures derived from blood transcriptomic data have often shown poor concordance across studies.^4,10^ These issues raise a critical question: can PD diagnostic methods based on blood-derived gene expression signatures be reliably replicated in independent datasets, and can performance be improved by controlling parameters that contribute to variability?

To address these questions, we systematically identified published blood transcriptomics-based gene signatures that had been developed to distinguish PD patients from controls. We then evaluated the performance of these signatures by using a large, independent dataset and by conducting a prospective trial designed to minimize potential confounders. Our findings show that while many signatures retain their statistically significant classification performance, the clinical applicability of these signatures remains limited— largely due to inter-individual variability, some of which can be ascribed to variance in genetic background.

## METHODS

### Data extraction from previous studies

We searched PubMed for articles (January 2015-May 2025), including “gene expression”, “RNA expression” or “transcriptomics”, plus “blood” and “Parkinson’s disease” and at least one of: “classification”, “AUC”, “ROC”, “sensitivity” or “specificity” (Fig. 1A). Of 91 articles identified, we included studies based on human mRNA reporting gene up- or downregulation.

**FIG. 1:**
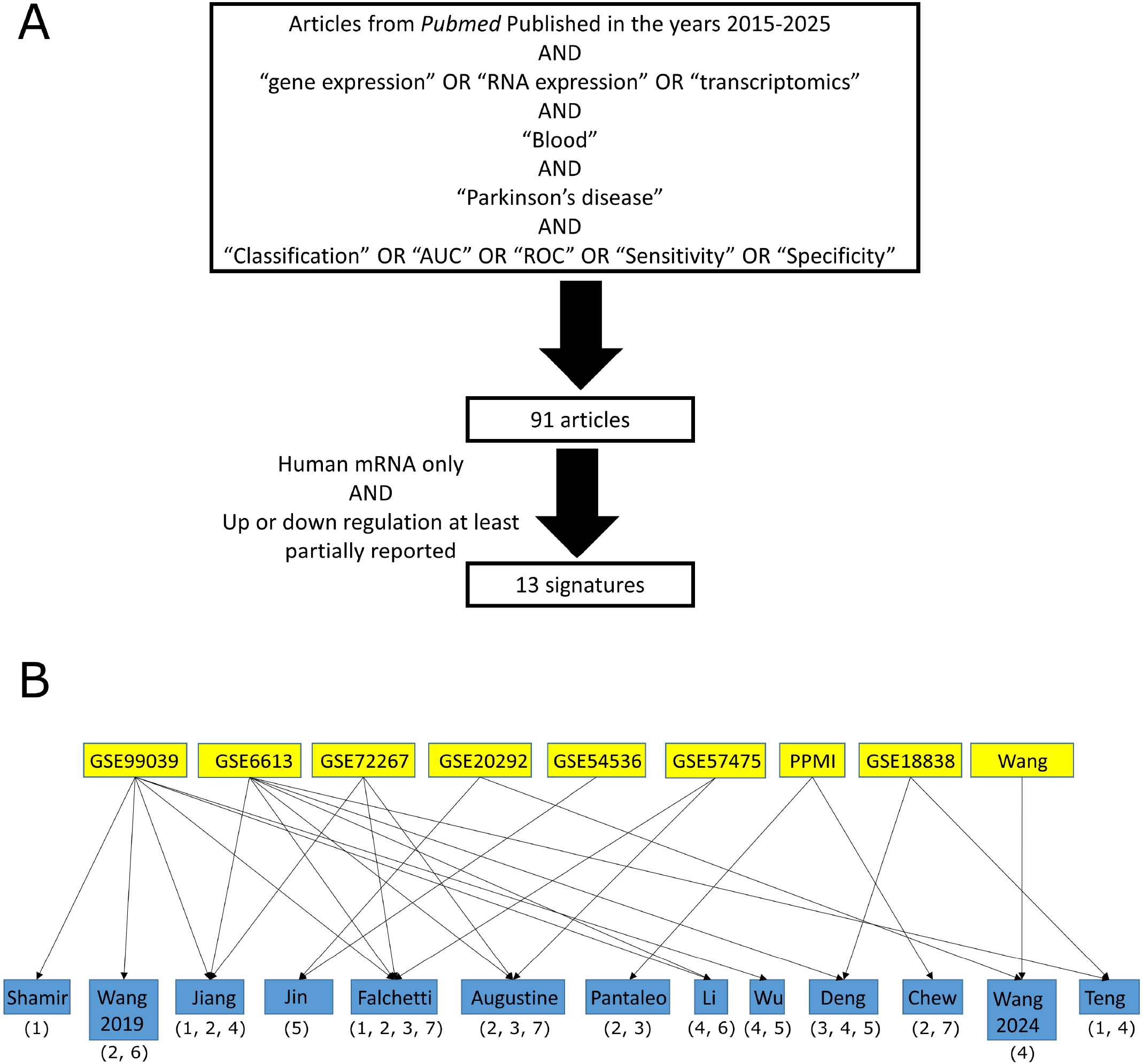
Reported signatures classifying PD and controls. A) Flowchart of the literature search, resulting in 13 studies. B) Original datasets (yellow boxes, upper panel) and the studies based on these datasets (blue boxes, lower panel) with the machine learning algorithms used for training the data. Digits represent the exact algorithm: 1) SVM (Support Vector Machine). 2) Random Forests. 3) Gradient Boosting. 4) LASSO (Least Absolute Shrinkage and Selection Operator). 5) Weighted Gene co-expression Network Analysis. 6) Methylation analysis. 7) Other type of analysis.

To enhance overlap between signatures, we focused on coding genes with reported regulation. In microarray data, if genes had multiple probes (as in Shamir et al.),^3^ we selected the probe with highest reported importance. We calculated random co-occurrence probability using 10,000 permutation tests. Gene enrichment was evaluated using Gene Set Enrichment analysis (GSEA)^11^ focusing on top 10 overlaps in KEGG Medicus gene sets, GO biological process and Human Phenotype Ontology, plus EnrichR^12^ focusing on WikiPathways 2024 Human, KEGG 2021 Human and GO Molecular Function 2025.

### Analyzing performance of signatures on external datasets

To classify participants based on the gene signatures, we introduced a simplified score that quantifies the relative expression of each gene signature in given sample. For RNA-sequencing datasets, we normalized the raw expression matrices by calculating counts per million (CPM) and for microarray datasets by quantile normalization. Normalization was followed by log2 transformation.

Given a normalized expression matrix M ∈ R^P×N^ measuring N signature genes across P participants, the score *s*_*j*_ of the j-th participant is given by:

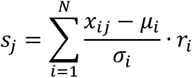

Here, for each signature gene, we standardized its expression across all samples in the cohort using the z-scores, and multiplied it by the regulation factor *r*_*i*_, representing up- or down-regulation of gene *x*_*i*_:

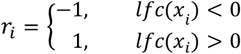

Where *lfc* = log-fold change between PD and control samples, reported in the original studies.

While weights based on log-fold change have been previously used in similar settings,^13^ we adopted a binary approach due to large variability between datasets (see results and previous studies^4^) and limited data on gene importance or log-fold change extractable from reported studies. Statistical analysis used 10,000 label (PD/control) permutations.

We validated our equal-weight approach by testing the performance of the two largest signatures (each from single dataset) using the original expression data it was based on. These signatures are particularly susceptible to bias from low-importance genes. For comparison, we repeated analysis after multiplying importance scores by +1 or -1 according to regulation direction. To test whether the number of differentially expressed genes in the consensus set was higher than expected by chance, we used the hypergeometric distribution separately for upregulated and downregulated genes, then combined the results using Fisher’s combined probability test.

Transcriptomics data was obtained from the Parkinson’s Progression Markers Initiative (PPMI).^14^ Data was extracted May 2025, including differential expression results and gene expression values for individual participants. We used the earliest sample from each participant. Demographic data included age on consent, sex, diagnosis and genetic status. Test set and source code used in Shamir et al. were downloaded from https://github.com/Shamir-Lab/GENEPARK. ^3^

To compare variability both between individuals and within individuals across two time points, we used mean pairwise distance (MPD), reflecting global similarity unaffected by sample number. MPD was calculated on normalized gene expression data by averaging absolute values of all relevant combinations within groups or differences before/after treatment for each individual. MPD was assessed for all signatures genes excluding those filtered for low counts in any of the groups. Group comparisons used Kruskal-Wallis test. General variability between individuals in different datasets was measured by MPD/mean and coefficient of variability (CV, standard deviation/mean). One-tailed Wilcoxon signed rank test assessed whether variability decreased after controlling environmental conditions.

### Prospective study to reduce expression variability

Participants were recruited at Hadassah Medical Center movement disorder clinic (June 2020-June 2022). All participants provided informed consent (IRB 0144-20-HMO). We included patients aged 50-80 diagnosed with PD following the MDS (Movement Disorders Society) criteria,^15^ naïve to symptomatic therapy (except MAO-B inhibitors), at clinical stage requiring symptomatic therapy initiation. Exclusion criteria were atypical PD, negative 6-[18F]fluoro-L-DOPA scan or prior neuroleptic use. Healthy controls were included after ruling out Parkinsonian signs. Baseline comparisons used Student’s t-test for age and Fisher’s exact test for sex.

PD patients and controls were tested following overnight fasting from caffeine-containing beverages and food. Upon arrival, participants were examined, motor part of the Unified Parkinson’s disease rating scale (UPDRS III) was calculated and 15 ml of peripheral blood for RNA-Seq was obtained. Then, some participants were treated with a tablet containing 125 mg of levodopa and 12.5 mg of carbidopa. Additional physical examination was performed and blood samples were obtained 150 minutes post-dose.

Blood samples were immediately preserved in RNA stabilization tubes (Tempus™) and stored in −80°C. Total RNA was extracted using the Tempus™ Spin RNA Isolation Kit, per manufacturer’s instructions. RNA concentration and purity were assessed using NanoDrop™ 2000 spectrophotometer (Thermo Fisher Scientific), and RNA integrity was evaluated using the Agilent 2100 Bioanalyzer (Agilent Technologies).

For each sample, 100 ng of total RNA was used to prepare RNA-seq libraries using the QuantSeq 3′ mRNA-Seq Library Prep Kit, per manufacturer’s protocol. To reduce globin mRNA-derived fragments, RS-Globin Block module was incorporated after first-strand cDNA synthesis. To eliminate amplification bias, standard Second Strand Synthesis Mix (SS1) was replaced with the UMI Second Strand Synthesis Mix (USS). Libraries from 41–44 blood samples were pooled and sequenced on Illumina NextSeq platform. Reads were aligned to the hg38 reference genome using STAR, and UMI-tools was used for UMI-based deduplication and generation of gene-level count matrices. Full reagents are in Table S1.

Data analysis used R (version 4.4.2; R Core Team, 2024) in RStudio environment and MATLAB R2024b (The MathWorks, Inc., Natick, MA). RNA-seq counts were processed using edgeR. Genes with expression levels below 1 CPM in fewer than three samples were filtered out, retaining 17,223 genes. Normalization used the trimmed mean of M-values (TMM) method followed by conversion to log2CPM. Correction for batch effect was performed using ComBat-seq. Differential expression to compare PD and control and time after levodopa intake was calculated using the limma R package.^16^ P-values were corrected via the Benjamini-Hochberg method. Cell-type deconvolution was performed using robust linear regression with non-negative constraints, using the LM22 signature matrix.^17^ The reference was scaled to match the distribution of TMM-normalized bulk data. Estimated proportions were normalized per sample, and neutrophil levels were compared across groups relative to calculated levels of B cells, T cells and natural killer cells.

## RESULTS

### Poor overlap between blood transcriptomics-based signatures

A literature search, based on our predefined criteria, identified 13 studies that classified PD and healthy control subjects based on blood-derived transcriptomic signatures (Fig. 1A, Table 1). Training and validation in these 13 studies were performed on 9 independent datasets (Fig. 1B). The median number of participants in each study was 205 PD patients (range: 19-400) and 189 controls (range: 19-369). 3/9 datasets included exclusively drug-naïve PD patients.

**Table 1:**
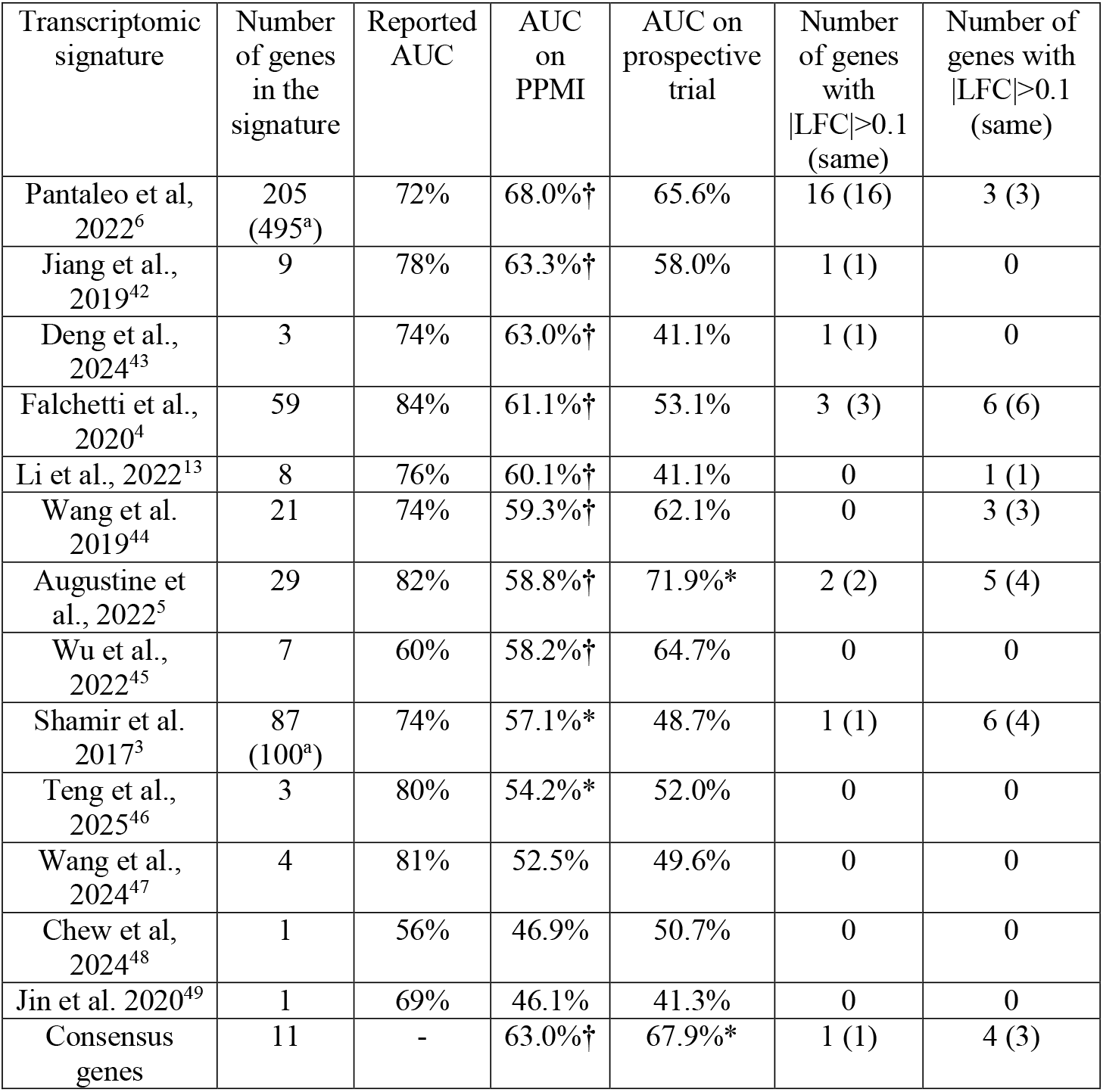
Classification performance and number of differentially expressed genes when applying the signatures on the PPMI and on our prospective data. * represents P<0.05; † represents P<0.001; AUC: area under the curve; Reported AUC: the maximum AUC reported in the original study on the test set. PPMI: Parkinson’s Progression Markers initiative; LFC: log-fold change; Same: number of genes with the same direction of regulation in both the original study and the external dataset. PM: post mortem. ^a^Originally reported including gene duplication, non-coding genes or genes with no reported regulation direction.

| Transcriptomic signature | Number of genes in the signature | Reported AUC | AUC on PPMI | AUC on prospective trial | Number of genes with $\vert \text{LFC}\vert > 0.1$ (same) | Number of genes with $\vert \text{LFC}\vert > 0.1$ (same) |
| --- | --- | --- | --- | --- | --- | --- |
| Pantaleo et al, 2022<sup>6</sup> | 205 (495<sup>a</sup>) | 72% | 68.0%† | 65.6% | 16 (16) | 3 (3) |
| Jiang et al., 2019<sup>42</sup> | 9 | 78% | 63.3%† | 58.0% | 1 (1) | 0 |
| Deng et al., 2024<sup>43</sup> | 3 | 74% | 63.0%† | 41.1% | 1 (1) | 0 |
| Falchetti et al., 2020<sup>4</sup> | 59 | 84% | 61.1%† | 53.1% | 3 (3) | 6 (6) |
| Li et al., 2022<sup>13</sup> | 8 | 76% | 60.1%† | 41.1% | 0 | 1 (1) |
| Wang et al. 2019<sup>44</sup> | 21 | 74% | 59.3%† | 62.1% | 0 | 3 (3) |
| Augustine et al., 2022<sup>5</sup> | 29 | 82% | 58.8%† | 71.9%* | 2 (2) | 5 (4) |
| Wu et al., 2022<sup>45</sup> | 7 | 60% | 58.2%† | 64.7% | 0 | 0 |
| Shamir et al. 2017<sup>3</sup> | 87 (100<sup>a</sup>) | 74% | 57.1%* | 48.7% | 1 (1) | 6 (4) |
| Teng et al., 2025<sup>46</sup> | 3 | 80% | 54.2%* | 52.0% | 0 | 0 |
| Wang et al., 2024<sup>47</sup> | 4 | 81% | 52.5% | 49.6% | 0 | 0 |
| Chew et al, 2024<sup>48</sup> | 1 | 56% | 46.9% | 50.7% | 0 | 0 |
| Jin et al. 2020<sup>49</sup> | 1 | 69% | 46.1% | 41.3% | 0 | 0 |
| Consensus genes | 11 | - | 63.0%† | 67.9%* | 1 (1) | 4 (3) |

Of the 9 datasets, transcriptomic profiling was performed using microarrays in 7 and RNA-sequencing in 2 (Fig. 1B). A median of 8 genes was identified across the 13 published signatures (range: 1-495), resulting in 411 unique genes at total after removing duplicates, non-coding transcripts and genes for which direction of regulation was not reported. 22/411 of these genes appeared in more than one signature but only half of them (11/411, 2.7%), i.e. consensus genes, originated from non-overlapping datasets (Table S2). While this overlap is small, it is unlikely to have occurred by chance (*P* < 0.001). None of these 11 consensus genes appeared in more than two independent signatures. Notably, all 11 consensus genes consistently showed the same direction of regulation (either up- or downregulated), indicating a degree of reproducibility.

Gene set Enrichment Analysis (GSEA) of the consensus genes revealed that 6/11 genes (*ABCA1, ABCG1, SLC25A20, SPON2, PTGDS* and *TREM1*) were related to lipid regulation (5/11 associated with response to lipids, corrected *P* = 0.012, 3/11 associated with intracellular lipid transport, corrected *P* = 0.004, Table S3). Similarly, analysis using EnrichR revealed the strongest enrichment in lipid-related pathways.

### Evaluation of classification signatures on the PPMI dataset

To assess the utility of each previously suggested signature on external datasets we implemented a simplified scoring system in which each gene was assigned a positive or negative weight based on its originally reported direction of regulation (up- or down-regulation respectively, see Methods). We first validated this approach by testing the two signatures with the largest number of genes both derived from a single dataset (Shamir et al., 2017, Pantaleo et al., 2022).^3,6^ Our scoring system successfully reproduced the ability of the original list of genes to distinguish PD from controls when applied to the relevant data sets (AUC 67% vs. reported 74% by Shamir et al. and AUC 68% vs. reported 72% by Pantaleo et al., Fig. S1). Both AUCs calculated by our scoring systems were significant (*P* = 0.014, *P* < 0.001 respectively). It is worth noting that for these two signatures, performances were similar when reported importance or effect sizes were used as weights (Shamir et al.: AUC 67.3%, *P* = 0.011; Pantaleo et al.: AUC 71.6%, *P* < 0.001) but since these metrics were only available in a subset of the studies, we opted to use equal weights to ensure consistency.

Applying our scoring system to all 13 gene signatures on the RNA-Seq based PPMI database revealed that 10/13 achieved statistically significant classification of PD vs. controls (*P* < 0.05, Table 1). Significant signatures calculated by our scoring system had median AUC of 59.7% (range: 54.2%-68.0%). This value was lower than originally reported, with a median decrease in AUC of 15.3% (range: 1.8%-25.8%). The three signatures that did not reach significance by our scoring system contained a smaller number of genes (1-4 genes in each signature). Of note, testing the classification performance of the 11 consensus genes achieved a relatively higher AUC of 63.0% (*P* < 0.001).

The lower classification performance calculated by our system, compared with the original studies, could be, at least partially, attributed to difference in reported gene expression between cohorts. While in the original studies most genes were considered to be differentially expressed as defined in the PPMI (log-fold change > 0.1 or < -0.1, e.g. 205/205 genes included from Pantaleo et al, 83/100 in Shamir et al., 59/59 in Falchetti et al.),^3,4,6^ in the PPMI database only 23/411 of the signature genes (5.84%) were differentially expressed. This limited overlap may be partially attributed to differences in platform technologies as most signatures derived from expression microarray data, while PPMI employed RNA-Seq.^10^ Of note, all 23 differently expressed signature genes exhibited the same direction of regulation as reported in the original reports (*P* = 0.001 for random occurrence).

### Assessing the classification performance on monogenic PD from the PPMI dataset

Since the etiology of PD is heterogeneous,^18^ we examined whether applying the suggested signatures on uniform genetic subgroups of PD from the PPMI database will increase the classification performance. For this analysis we included 304 patients with negative genetic tests (idiopathic, iPD), 215 patients positive for pathogenic variants in *LRRK2* (*LRRK2*-PD), 103 patients positive for pathogenic variants in *GBA1* (*GBA*-PD) and 161 healthy controls with a negative genetic panel. We analyzed the 10/13 signatures that exhibited a statistically significant classification performance.

Limiting the classification to only *LRRK2*-PD vs. controls improved performance in 7/10 signatures, although this overall improvement was not statistically significant relative to the ability of these signatures to distinguish iPD from controls (*LRRK2*-PD vs. controls: median AUC 61.3%, range 51.1-65.2%; iPD vs. controls: median AUC 57.1%, range 54.3-68.1%; *P* = 0.160, Fig. S2). In *GBA*-PD, 9/10 signatures did better in distinguishing PD from controls (*GBA*-PD vs. controls median AUC 65.4%, range: 54.4-72.7%; iPD vs. controls: median AUC 57.1%, range: 54.3-68.1%, *P* = 0.006). Improved performance in distinguishing *GBA*-PD from controls was also noticed in the signature that was based on data from which monogenic PD were excluded.^6^ With the 11 consensus genes, AUC was 71.2% in *GBA*-PD vs. controls as comparison to 63.8% in iPD vs. controls.

Although this study does not focus on asymptomatic carriers, it is noteworthy that 8/11 of the consensus genes are differentially expressed between *GBA*-PD and unaffected *GBA*1 mutations carriers, and 7/11 are differentially expressed between *LRRK2*-PD and unaffected *LRRK2* mutations carriers (*P* < 0.001 for both comparisons, hypergeometric test was applied to overcome effect secondary to different sample sizes). The direction of expression change in all of these genes (comparing monogenic PD to non-manifesting carriers) was consistent with that observed in the original studies comparing PD to healthy controls (Fig. S3).

### Source of gene expression variability in the PPMI dataset

Previous studies have suggested that the classification performance of blood transcriptomics-based signatures is limited, in part, by the variability of gene expression.^10^ This variability may rise from either intra-individual differences (within the same individual) documented in repeated sampling or inter-individual differences (between different individuals), and in both cases can be affected by environmental conditions. While most environmental factors are difficult to control or quantify, the use of anti-parkinsonian therapy represents a measurable and clinically relevant variable. Indeed, the use of dopamine-replacement therapy has been hypothesized to influence gene expression independently of PD presence, potentially confounding transcriptomic signatures.^19,20^ Notably, 11/13 gene signatures analyzed in our study were derived from cohorts in which at least part of the participants received dopaminergic therapy.

In order to assess the potential contribution of intra- and inter-individual differences and exposure to dopaminergic therapy to variability of gene expression, we computed the mean pairwise distance (MPD) between individuals with PD and between healthy controls prior to treatment initiation. We also calculated the average gene expression change within each PD patient over the year before and after starting therapy, and for controls over equivalent time intervals.

Among all signature genes with sufficient counts (282/411) MPD values were similar between PD patients and controls (*P* = 0.951; Fig. 2A), suggesting comparable variability between individuals in both groups. As expected, MPD values within individuals across two time points were significantly lower (*P* < 0.001 for both PD and controls), reflecting lower variability within each individual across different time-points, even with anti-parkinsonian therapy. Importantly, the gene expression change following the initiation of anti-parkinsonian therapy in PD patients did not exceed the natural variability observed across time points in untreated controls (*P* = 0.960). These findings suggest that the dominant source of variability in blood gene expression lies between individuals, rather than within individuals over time, and that long-term dopaminergic therapy does not substantially increase variability in gene expression profiles.

**FIG. 2:**
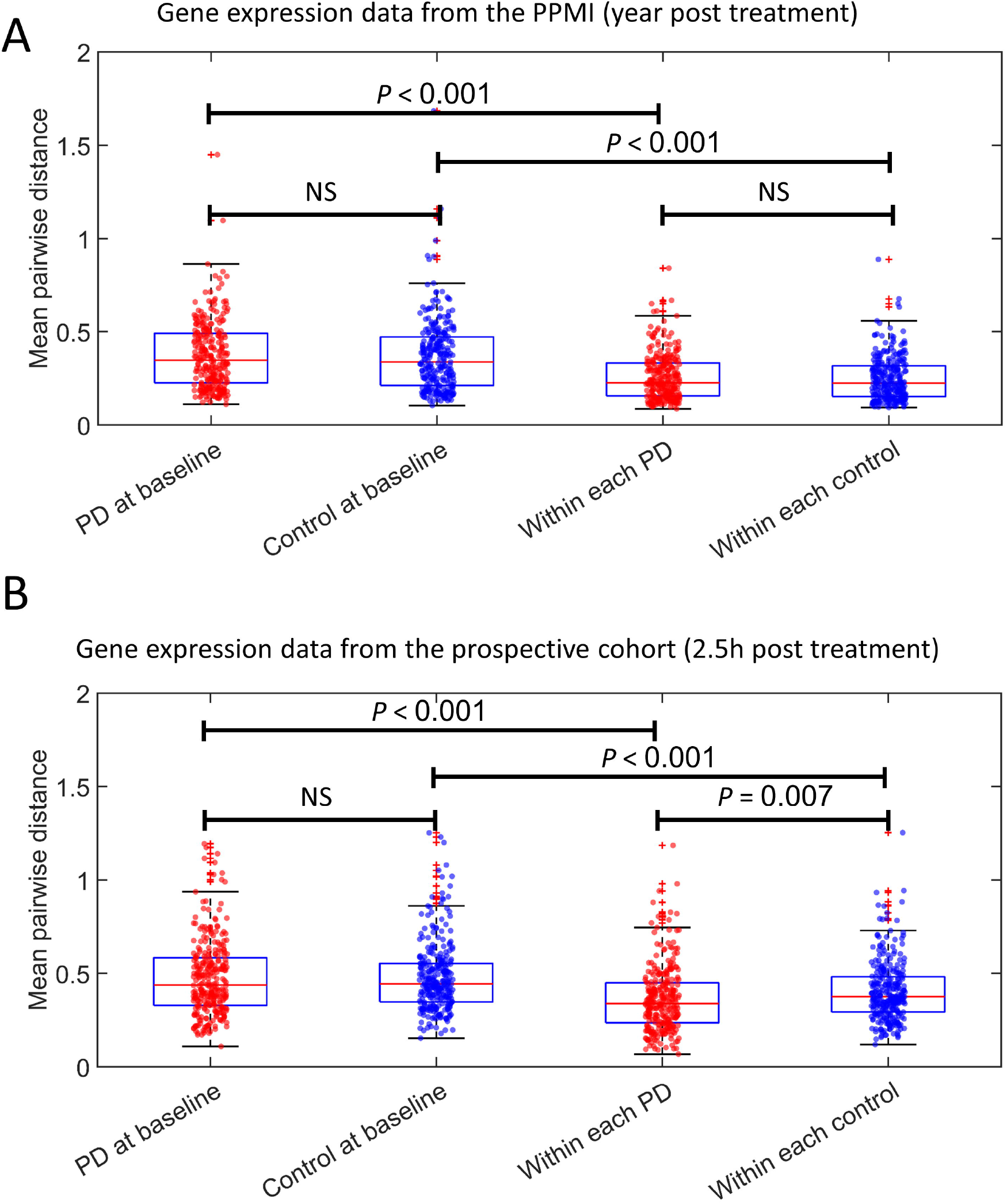
Mean pairwise distance (MPD) of gene expression values from blood RNA-seq across signature genes. A) MPD of data from the PPMI of 282 genes, calculated between treatment-naïve PD and controls and across two time points—in the year before and in the year after initiating anti-parkinsonian therapy for PD patients, and at matched time intervals for controls. MPD was significantly lower within individuals than between individuals (PD at baseline: median 0.35 [0.11-1.45], within each PD: median 0.23 [0.09-0.84], *P* < 0.001; Controls at baseline: median 0.34 [0.10-1.68], within each control: median 0.22 [0.09-0.89], *P* < 0.001). No significant difference was observed between the treatment-related change in PD patients and the natural variation in controls (*P* = 0.960), suggesting that long-term anti-parkinsonian therapy does not substantially increase gene expression variability. B) MPD of data from an independent prospective cohort of 16 drug-naïve individuals (6 PD, 10 controls) of 299 genes. MPD was calculated among participants before receiving treatment (baseline) and within each participant 150 minutes after receiving 125/12.5 mg of levodopa/carbidopa. In both PD patients and controls, variability within individuals across time points was significantly lower than variability between individuals at baseline (PD at baseline: median 0.44 [0.11-1.19], within each PD: median 0.34 [0.07-1.18], *P* < 0.001; Controls at baseline: median 0.44 [0.15-1.25], within each control: median 0.38 [0.12-1.25], *P* < 0.001). The treatment-induced change was smaller in PD than in controls (*P* = 0.007), indicating a modest but consistent within-subject response. PPMI: Parkinson’s Progression Markers initiative. NS: non-significant. PD: Parkinson’s disease.

### Intervention study attempting to reduce gene-expression variability

We hypothesized that controlling environmental factors and reducing clinical and demographic differences between individuals, would reduce the variability of gene expression and improve the performance of the signatures. In addition, this may reduce some inconsistencies observed in gene regulation between cohorts previously described also for other blood transcriptomics signatures.^10^ Towards this aim, we performed a prospective clinical trial (30 participants, 16 patients with PD and 14 healthy controls, see Table S4 for demographic characteristics). We minimized factors known to affect blood transcriptome by recruiting only patients at the same clinical stage, i.e. patients naïve to symptomatic therapy required to initiate treatment.^21^, narrowing the age range (50-80 years),^22,23^ sampling blood at the same time of the day (7-8 AM) ^24^ after a night fast,^25^ and requesting participants to avoid caffeinated beverages for at least 12 hours.^26^ The neutrophil-to-lymphocyte ratio, a potential confounder,^27^ reported to differentiate PD from controls,^28^ was later calculated, and showed no significant difference between groups (PD: 1.55±0.58, Controls: 1.27±0.34, *P* = 0.118). All participant demographics are provided in Table S4.

In contrast to our hypothesis, the expression variability between individuals did not decrease neither in the PD (MPD/mean: *P* = 0.813; Coefficient of variation: *P* = 1.000) nor in the control groups (MPD/mean: *P* = 0.927; Coefficient of variation: *P* = 1.000). Similar to our results that were based on the PPMI dataset, also here only 20/411 (4.87%) were differentially expressed in PD vs. controls, 18/20 (90.0%) of them with the same direction of regulation (Table 1). As neither the variability nor the rate of differential expression improved in our cohort, classification performances were mostly similar to the results from the PPMI (in our cohort: median AUC 52.0%, range: 41.1%-72.9%; PPMI: median AUC 58.8%, range: 46.1%-68.0%; median difference between AUCs 3.0%, *P* = 0.216; Table 1, Fig. S4).

To examine whether levodopa induces short-term changes, in either PD or healthy individuals, 16 of the participants (6 with PD and 10 healthy controls) received a single oral dose of 125/12.5 mg levodopa/carbidopa for the first time in their life, and peripheral blood was collected for RNA sequencing 150 minutes later. Consistent with the long-term findings from the PPMI cohort, gene expression variability, measured by MPD, was significantly higher between individuals than within individuals (i.e., before and after levodopa intake) at baseline (*P* < 0.001 for both PD and controls; Fig. 2B). Interestingly, MPD values after levodopa were smaller in PD patients (*P* = 0.007), suggesting a modest differential transcriptional short-term response to dopaminergic stimulation.

## DISCUSSION

In this study, we identified a small but significant overlap (11/411) of consensus genes across previously proposed transcriptomic signatures designed to distinguish PD patients from healthy controls. While we validated the ability of 10/13 signatures to recognize PD, AUCs were overall low when applied to external datasets (median AUC 59.7%). Improvement, to a certain degree, was observed when exclusively analyzing *GBA*-PD individuals. Inter-individual gene expression variability, a major limitation in signature performance, was not reduced by prospective attempts to optimize clinical, demographic and environmental factors of participants. We concluded that biological differences between individuals, rather than environmental or treatment effects, are the dominant source of gene expression variation and limit the diagnostic value of blood-based expression signatures.

It is now well established that PD-related molecular changes are not confined to the brain. PD-related transcriptomic alterations have been observed in the skin,^29^ gastrointestinal system,^30^ and as was confirmed here, also in blood. These blood transcriptomic changes were shown to mirror RNA changes observed postmortem in the striatum, underscoring their potential relevance to disease biology.^31^ While the clinical utility of blood transcriptomic changes remain limited (see below), identifying reproducible molecular alterations is essential for deepening our understanding of PD pathophysiology. In this regard, it is noteworthy that 6/11 of the consensus genes are involved in lipid transport or lipid-regulatory pathways, aligning with prior reports of PD-related disruptions in lipid metabolism and its connection to alpha-synuclein pathology.^32^

Although the ability of blood transcriptomics-based signatures to distinguish patients with PD from controls, as was reproduced here, is overall statistically significant, their performance is not clinically useful. Among the 10 of 13 signatures that retained statistical significance in our analysis, the highest AUC was only 68.0% which is considered to be an inadequate classifier.^33^ While our use of an equal-weighted scoring approach may have contributed to reduced performance, even in the original publications none of the signatures exceeded AUC of 85%, which is considered good^33^ but by itself does not imply clinical utility^34^ and falls short of the clinical diagnosis of PD by movement disorders specialists (AUC 92.5%).^35^

One plausible explanation for the limited performance and low concordance among gene signatures is the genetic and pathological heterogeneity of PD.^18^ Most notably, gene signatures tended to perform better in patients with *GBA*-PD, an accentuated form of the disease,^36^ that is uniformly associated with alpha-synuclein pathology.^37^ Conversely, when analyzing data for *LRRK2*-PD, which is associated with a more heterogeneous pathology,^38^ there was not a significant improvement in signature performances. This supports the notion that genetic stratification may improve biomarker consistency and interpretability.

A key challenge in developing blood transcriptomics-based classifiers is disentangling disease-specific transcriptional signatures from confounding variables. Contrary to prior assumptions,^19^ our findings suggest that anti-parkinsonian therapy initiated within the year preceding sampling does not substantially contribute to gene expression variability. Previous studies noted several other potential confounders - such as age, time of day, fasting status, caffeine intake and proximity to levodopa administration.^20,22–26^ However, even in our prospective cohort, where such factors were carefully minimized, variability between individuals persisted and classification accuracy did not improve. Future studies may benefit from stricter methodological standardization, including adjustments for blood cell sub-populations, to enhance reproducibility.^39^

Finally, with the advent of highly sensitive and specific diagnostic methods such as α-synuclein seed amplification assays,^40^ the rationale for pursuing a universal transcriptomic signature of Parkinson’s disease becomes increasingly questionable. Instead, leveraging the unique gene expression patterns of individual patients may enable the development of divergent biomarkers that capture personalized disease trajectories.In conclusion, we identified a modest but statistically significant overlap among published blood transcriptomics-based gene signatures for PD, alongside substantial variability between individuals that limits their clinical applicability. While future methods—including those leveraging blood-derived mRNA from single-cells or non-coding RNAs^41^ may yield more robust biomarkers, current evidence suggests that pathology-driven approaches, such as alpha-synuclein seed amplification assays, offer superior clinical potential and reliability.^40^ Rather than pursuing a universal PD transcriptomic signature, efforts should leverage RNA variability to develop personalized biomarkers that reflect individual disease trajectories.

## AUTHORS’ ROLES

(1) Research project: A. Conception, B. Organization, C. Execution; (2) Statistical Analysis: A. Design, B. Execution, C. Review and Critique; (3) Manuscript: A. Writing of the first draft, B. Review and Critique.

RD: 1A, 1B, 1C, 2A, 2B, 2C, 3A, 3B

SD: 1A, 1B, 1C, 3B

HT: 1A, 1B, 1C, 3B

MG: 2A, 2B, 2C, 3A, 3B

SS: 1A, 1B, 1C, 3B

SH: 1A, 1B, 1C, 3B

AC: 1A, 1B, 1C, 3A, 3B

TH: 1A, 1B, 1C, 2A, 2B, 2C, 3A, 3B

DA: 1A, 1B, 1C, 2A, 2B, 2C, 3A, 3B

## FINANCIAL DISCLOSURES (for the preceding 12 months)

The authors have no financial interest to disclose.

## DATA AVAILABILITY STATEMENT

Data used in the preparation of this article were obtained on [2025-05-01] from the Parkinson’s Progression Markers Initiative (PPMI) database (https://www.ppmi-info.org/access-data-specimens/download-data), RRID:SCR_006431. For up-to-date information on the study, visit http://www.ppmi-info.org.

Data from the prospective cohort is available upon request.

## ACKNOWLEDGEMENT

RD received support from the Israeli Center for Addictions (ICA). RD and DA received support from Berel and Agnes Ginges family foundation. Initial support for the project was awarded from the Israel Ministry of Health to DA and AC (Grant 3-17303). PPMI – a public-private partnership – is funded by the Michael J. Fox Foundation for Parkinson’s Research and funding partners, including 4D Pharma, Abbvie, AcureX, Allergan, Amathus Therapeutics, Aligning Science Across Parkinson’s, AskBio, Avid Radiopharmaceuticals, BIAL, BioArctic, Biogen, Biohaven, BioLegend, BlueRock Therapeutics, Bristol-Myers Squibb, Calico Labs, Capsida Biotherapeutics, Celgene, Cerevel Therapeutics, Coave Therapeutics, DaCapo Brainscience, Denali, Edmond J. Safra Foundation, Eli Lilly, Gain Therapeutics, GE HealthCare, Genentech, GSK, Golub Capital, Handl Therapeutics, Insitro, Jazz Pharmaceuticals, Johnson & Johnson Innovative Medicine, Lundbeck, Merck, Meso Scale Discovery, Mission Therapeutics, Neurocrine Biosciences, Neuron23, Neuropore, Pfizer, Piramal, Prevail Therapeutics, Roche, Sanofi, Servier, Sun Pharma Advanced Research Company, Takeda, Teva, UCB, Vanqua Bio, Verily, Voyager Therapeutics, the Weston Family Foundation and Yumanity Therapeutics.

