## Supplemental for "The Quest for a Universal Parkinson’s Transcriptomics Signature is Derailed by Inherent Variability between Patients"

**SUPPLEMENTARY FIGURES**

**FIG S1:**

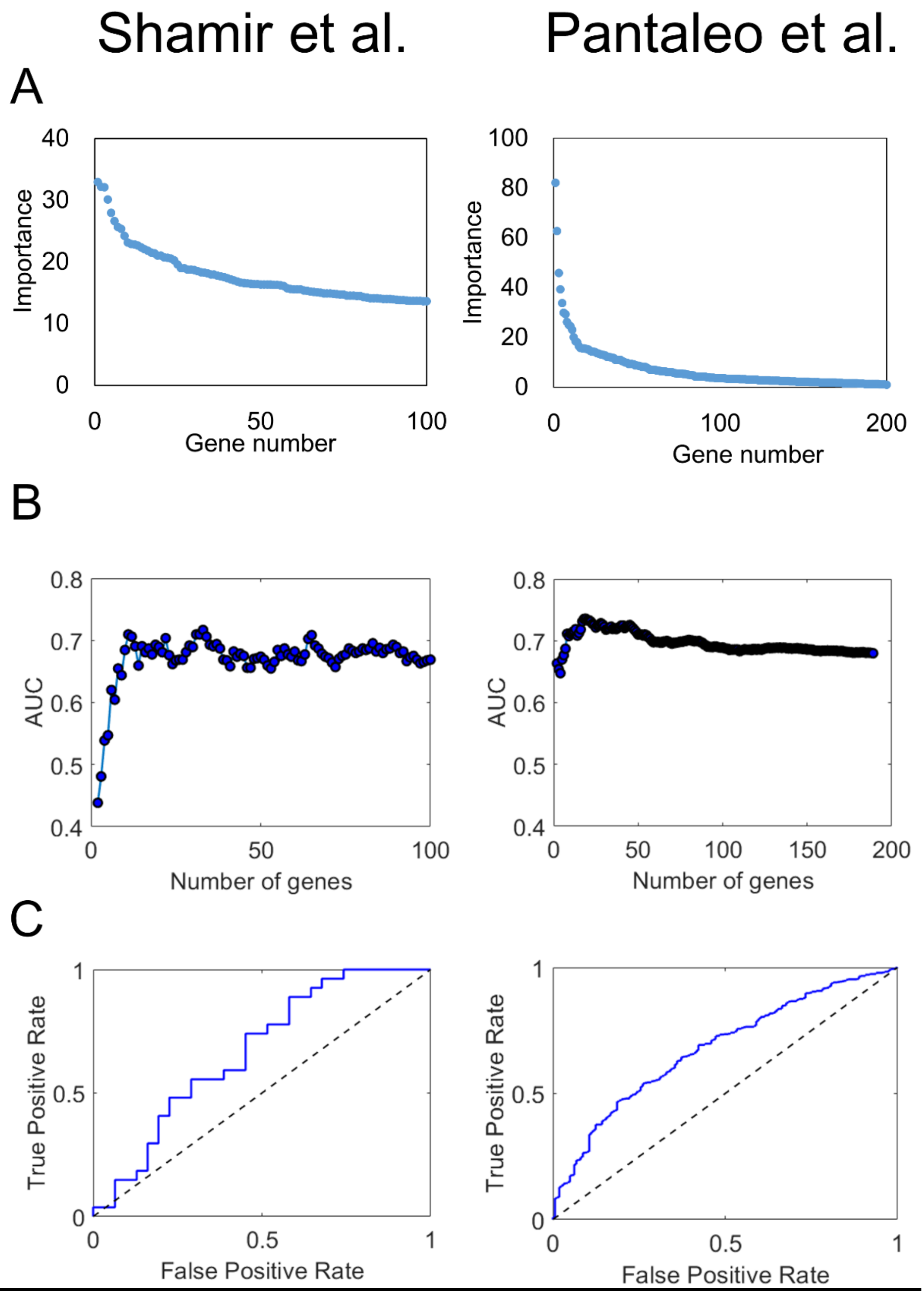


FIG S1: Validation of our scoring method using the signature from Shamir et al. on the test set from the study (left panel) and Pantaleo et al. on data from the PPMI on which it was based (right panel, only genes with reported upregulation or downregulation). A) Importance of each gene from the signature in a descending order, as reported by the original study. B) AUC of the signature when applying the first N genes. In both signatures there is an initial steep rise in AUC after the first few genes, reflecting the genes with highest importance. This peak is similar to the AUC in the original study (Shamir et al.: 72% vs. 74% originally reported; Pantaleo et al.: 74% vs. 72% originally reported). The peak AUC is followed by a gradual decrease, reflecting the addition of lower-importance genes that undermine the weight of the first genes. C) ROC curve of the signatures after applying all genes using our method, yielding a significant classification performance (AUC 67%, $P = 0.011$ for Shamir et al.; AUC 68%, $P < 0.001$ for Pantaleo et al.), despite being somewhat lower than the original AUC reported and the peak AUC. PPMI: Parkinson's Progression Marker Initiative. AUC: Area under the curve. ROC: receiver operator characteristics.

**FIG S2:**

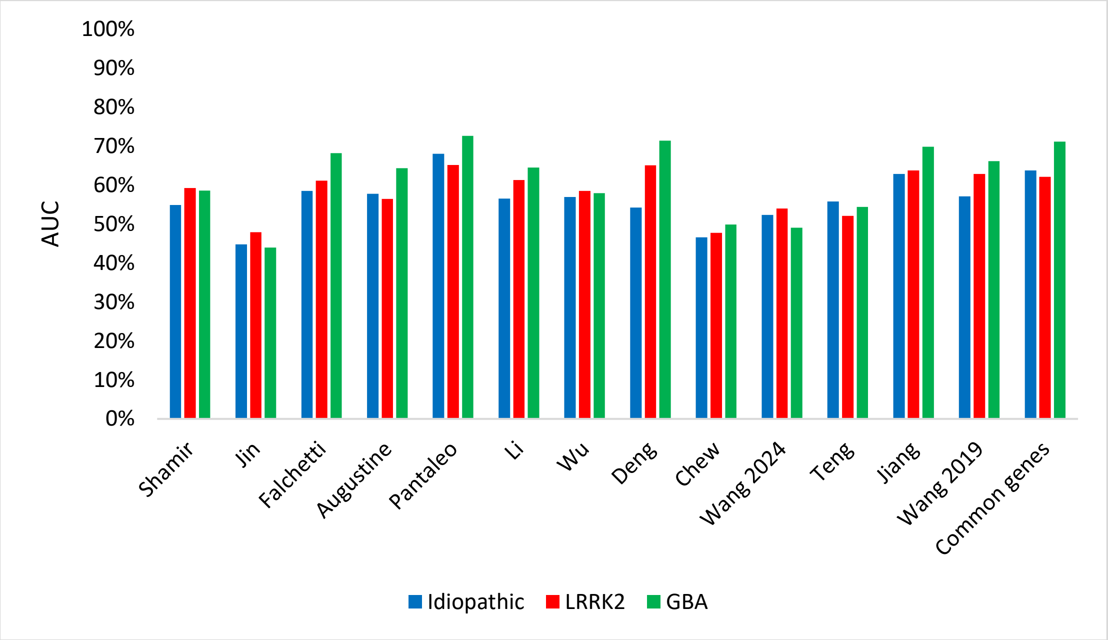


FIG. S2: Classification performance of different signatures applied on different genetic subpopulations of PD from the PPMI database vs. non-carrier healthy controls. AUC: Area under the Curve.

**FIG S3:**

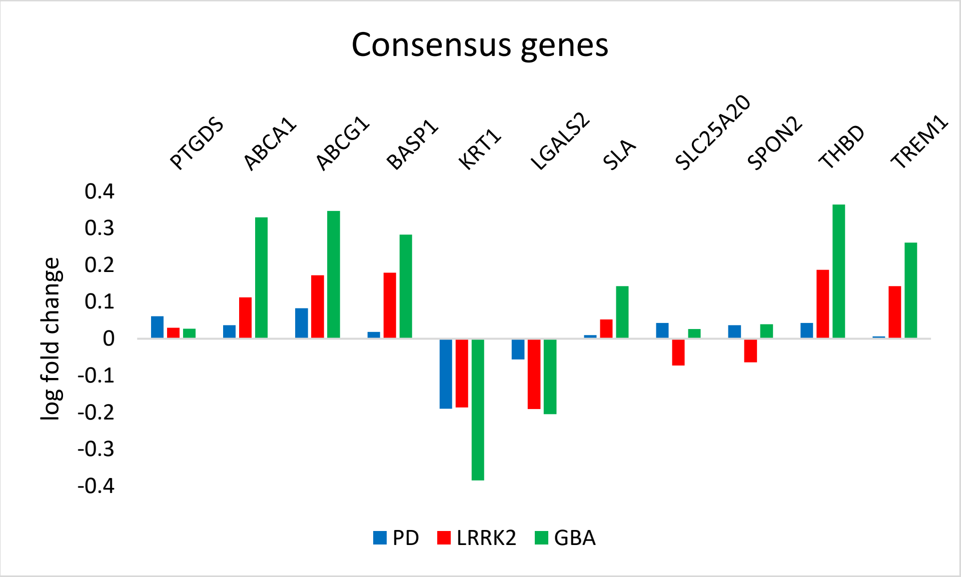


FIG. S3: Comparison of differential expression of PD patients from the PPMI in the 11 consensus genes according to genetic etiology. Blue: general comparison to PD vs. healthy controls. Orange: *LRRK2*-PD vs. non-manifesting carriers of *LRRK2*. Grey: *GBA*-PD vs. non-manifesting carriers of *GBA*. Values bellow -0.1 or above 0.1 are considered differentially expressed.

**FIG. S4:**

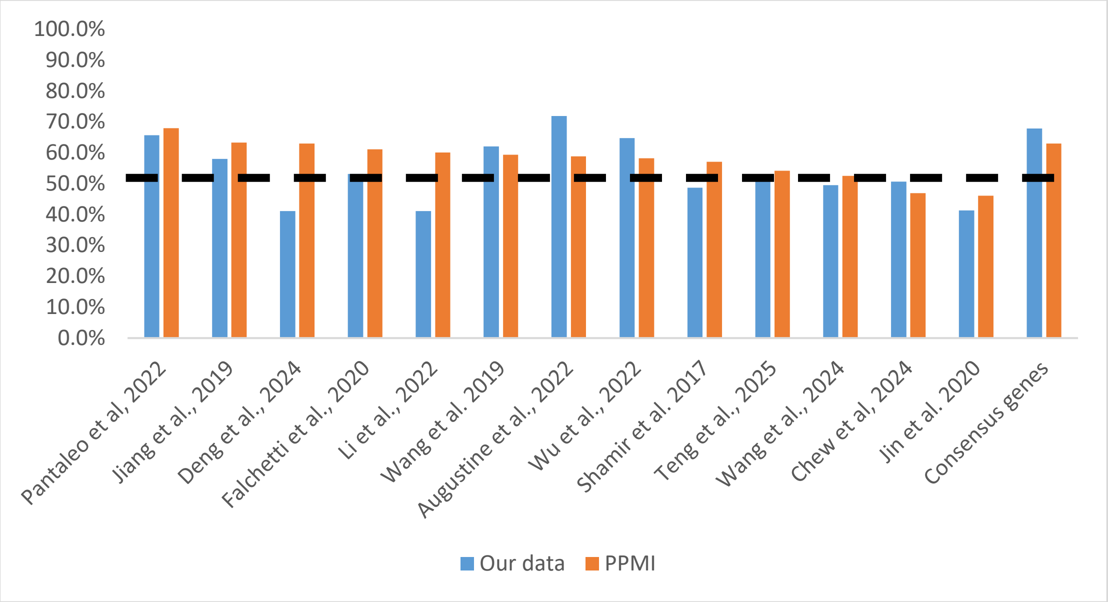


FIG. S3: Comparison of classification performances of the different signatures on our prospective cohort (blue) and data from the PPMI (orange).

### SUPPLEMENTARY TABLES

Table S1: Reagents used in the RNA-Seq analysis

| **reagent** | **Catalog number** | **company** |
|---|---|---|
| Tempus™ Blood RNA Tube | AB-4342792 | Invitrogen |
| Tempus™ Spin RNA Isolation Kit | AB-4380204 | Invitrogen |
| qScript cDNA Synthesis Kit | 95047-100 | Quantabio |
| QuantSeq 3‘ mRNA-Seq Library Prep Kit (FWD) for Illumina, 96 preps | LX-015.96 | Lexogen |
| RS-Globin Block, Homo sapiens,96 rxn | LX-070.96 | Lexogen |
| UMI Second Strand Synthesis Module for QuantSeq FWD (Illumina, Read 1), 96 rxn | LX-081.96 | Lexogen |

Table S2: Consensus genes – occurring more than once in independent signatures (1: upregulated, -1: downregulated)

| **Gene** | **Regulation** | **Number of occurrences** |
|---|---|---|
| *PTGDS* | 1 | 2 |
| *ABCA1* | 1 | 2 |
| *ABCG1* | 1 | 2 |
| *BASP1* | 1 | 2 |
| *KRT1* | -1 | 2 |
| *LGALS2* | -1 | 2 |
| *SLA* | 1 | 2 |
| *SLC25A20* | 1 | 2 |
| *SPON2* | 1 | 2 |
| *THBD* | 1 | 2 |
| *TREM1* | 1 | 2 |

Table S3: Gene set overlap in the consensus genes from GSEA and the relevant gene sets.

| Entrez Gene Id | Gene Symbol | Gene set |
|---|---|---|
| 19 | ***ABCA1*** | **1, 2, 3, 5, 6, 7, 8, 9, 10** |
| 9619 | ***ABCG1*** | **1, 2, 3, 5, 6, 7, 8, 9, 10** |
| 788 | ***SLC25A20*** | **1** |
| 10417 | ***SPON2*** | **2, 4** |
| 5730 | ***PTGDS*** | **2, 4** |

| | | |
|---|---|---|
| 7056 | *THBD* | 2 |
| 54210 | ***TREM1*** | **4** |
| 3848 | *KRT1* | |
| 10409 | *BASP1* | |
| 3957 | *LGALS2* | |
| 6503 | *SLA* | |

| Reference number | Gene Set Name | p-value | FDR q-value |
|---|---|---|---|
| **1** | **GOBP Intracellular Lipid Transport** | 2.50E-07 | 3.50E-03 |
| **2** | **GOBP Response To Lipid** | 2.11E-06 | 1.20E-02 |
| **3** | **GOBP Negative Regulation Of Cholesterol Storage** | 3.21E-06 | 1.20E-02 |
| 4 | GOBP Myeloid Leukocyte Mediated Immunity | 3.43E-06 | 1.20E-02 |
| 5 | GOBP Negative Regulation Of Macrophage Derived Foam Cell Differentiation | 5.30E-06 | 1.24E-02 |
| **6** | **GOBP Phospholipid Efflux** | 5.30E-06 | 1.24E-02 |
| **7** | **GOBP Phospholipid Homeostasis** | 7.92E-06 | 1.58E-02 |
| **8** | **GOBP Regulation Of Cholesterol Storage** | 9.96E-06 | 1.74E-02 |
| **9** | **GOBP Reverse Cholesterol Transport** | 1.22E-05 | 1.88E-02 |
| **10** | **GOBP Negative Regulation Of Lipid Storage** | 1.34E-05 | 1.88E-02 |

GSEA: Gene set Enrichment Analysis. GOBP: Gene Ontology Biological Process. Bold: related to lipid metabolism or regulation.

Table S4: Demographic details of participants from the prospective cohort

| **Our data** | **PD (N=16)** | **controls (N=14)** | **p-value** |
|---|---|---|---|
| Age (mean, STD, range) | 66.8± 6.9 [51-75] | 62.9 ± 8.3 [50-78] | 0.179 (t-test) |
| Sex (F/M) | 7/9 | 9/5 | 0.299 (Fisher's exact test) |
| UPDRS-3 (median, IQR range) | 22 ± 13 (13-37) | NA | NA |
| Disease duration (median, IQR range) | 2 ± 2 (0-5) | NA | NA |
| *LRRK2*+ | 2 (12.5%) | NA | NA |
| *GBA*+ | 0 (0%) | NA | NA |

PD: Parkinson's disease. STD: Standard deviation. F: Females. M: Males. UPDRS-3: Unified Parkinson's disease rating scale part 3. IQR: Interquartile range. NA: non-applicable.
